# Inter-slice motion correction using spatiotemporal interpolation for functional magnetic resonance imaging of the moving fetus

**DOI:** 10.1101/204404

**Authors:** Wonsang You, Catherine Limperopoulos

## Abstract

Fetal motion continues to be one of the major artifacts in in-utero functional MRI; interestingly few methods have been developed to address fetal motion correction. In this study, we propose a robust method for motion correction in fetal fMRI by which both inter-slice and inter-volume motion artifacts are jointly corrected. To accomplish this, an original volume is temporally split into odd and even slices, and then voxel intensities are spatially and temporally interpolated in the process of image registration. Our experimental data demonstrate that our method was more effective in correcting fetal motion artifact compared to traditional motion correction methods.

## Introduction

Fetal movement seriously deteriorates the quality of in-utero functional MR images by causing geometric mismatch between volumes. In addition, it gives rise to geometric distortion of intensities between subsequent slices which are acquired sequentially. A few methods have been suggested to correct interslice motion as well as inter-volume motion(1–4). The objective of this study was to propose a robust method where spatial distortion of structure and intensities which is present between slices is effectively corrected considering temporal variation of voxel intensities.

## Methods

The proposed method for motion correction is organized into five steps: temporal expansion, spatial interpolation, image registration, temporal interpolation, and temporal compression as depicted in Figure 1. First, an original 4D sequence of length L with time interval Δt(=TR) is temporally expanded into a sequence of length 2L with interval Δt/2 by alternating odd and even slices as it was acquired with interleaved scan. The missing slices in each expanded volume are filled using shape-preserving piecewise cubic interpolation over slices. Both inter-slice and inter-volume motion artifacts are then simultaneously corrected by registering each expanded volume into a given template image using either rigid or non-rigid transformation. Voxel-wise temporal interpolation between time points is applied based on linear polynomial curve fitting to correct the temporal variation between odd and even slices. Finally, the temporal resolution of Δt/2 is reduced to the original interval Δt by taking just odd volumes.

**Figure 1.**
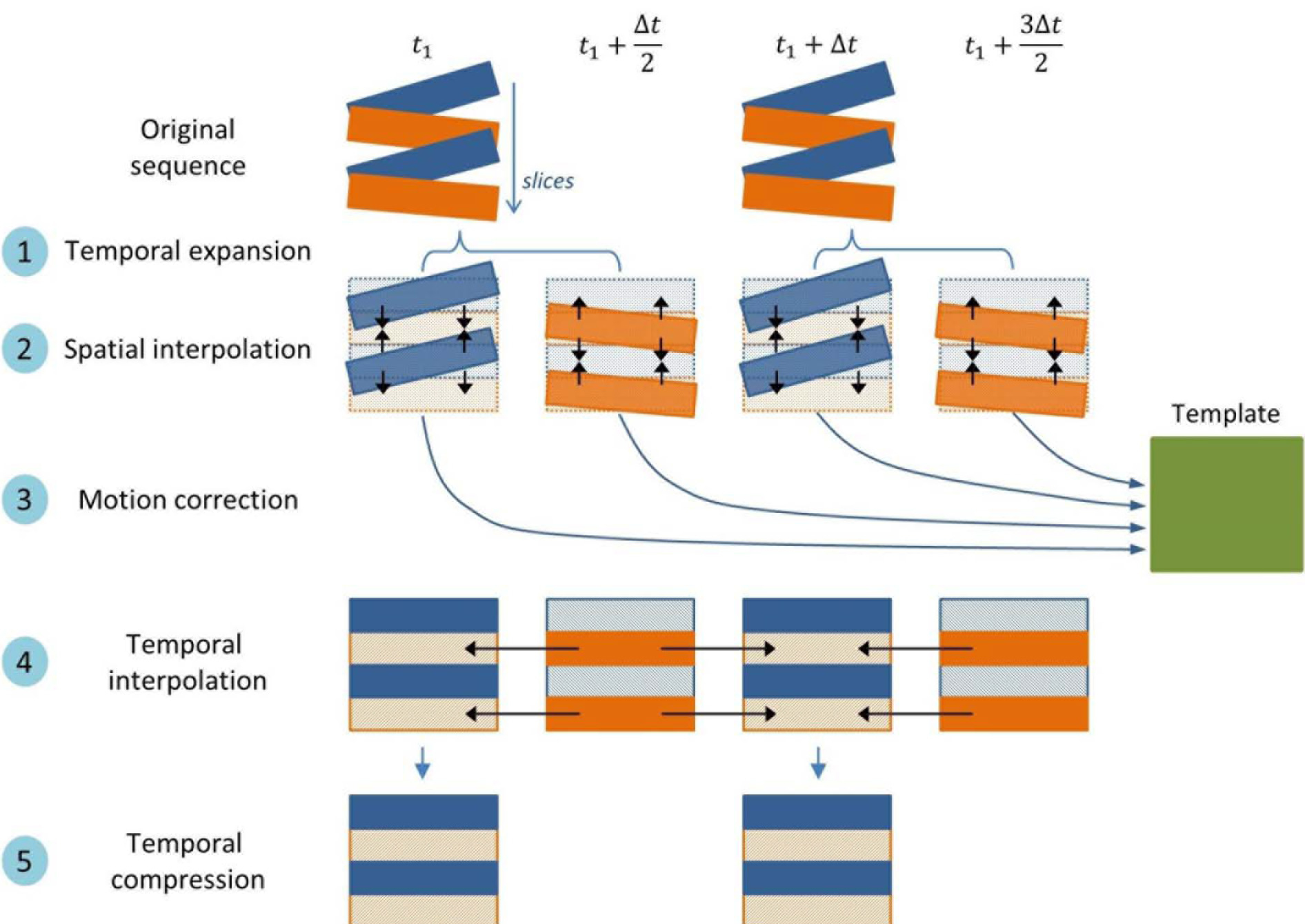
The overall process of spatiotemporal motion correction.

To evaluate the performance of the proposed method, four healthy pregnant women at 28-37 weeks gestation were examined with BOLD fMRI on a 1.5T MR scanner (GE). A volumetric MR image sequence including the whole placenta was acquired for each subject during 4-minute maternal hyperoxia with imaging parameters TE=1s, TR=2s, flip angle=90°, voxel size 3.28×3.28×8 mm^3^, and slice gap 2 mm. Rigid image registration was applied for simulated data using FLIRT (as a part of FSL)(5) while non-rigid registration was applied for in-utero fMRI data using Elastix(6). Structural similarity index (SSIM), which was related to correlation in intensities and structure between volume and template, was then computed as a metric to assess the performance of motion correction (7).

## Results

Figure 2 illustrates that the spatial mismatch and intensity distortion between slices caused by motion artifact were effectively eliminated from an fMRI volume of the placenta. As listed in Table 1, The SSIM score improved by 4.53% on average after both spatial and temporal interpolations were jointly applied. On the other hand, the SSIM score negligibly increased by only 0.43% on average after only spatial interpolation was applied as proposed in (8).

**Figure 2.**
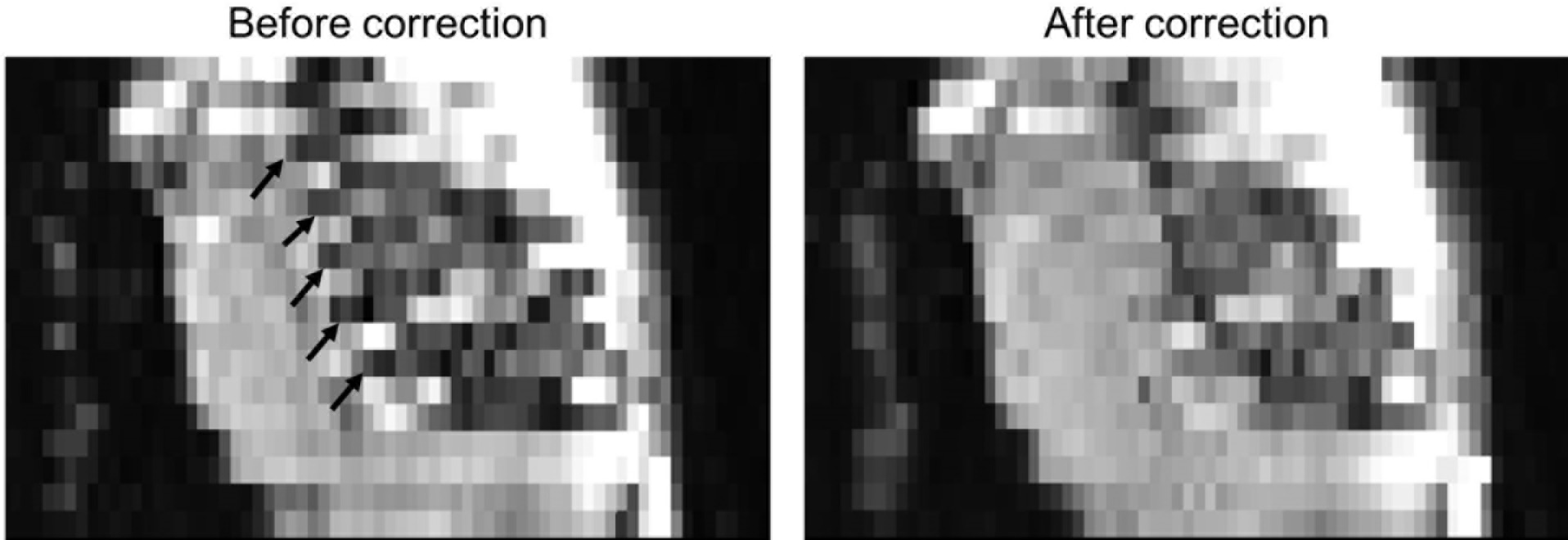
The effects of the proposed motion correction method on in-utero fMRI data. The original volume was seriously perturbed by inter-slice motion artifacts (see arrows). After applying the proposed method, Intensity distortion caused by inter-slice motion

**Table 1.** The SSIM scores before and after motion correction in each subject

|  | SSIM |  |  |
| --- | --- | --- | --- |
|  | Original | Spatial interpolation only | Spatial + temporal interpolation |
| Subject 1 | 0.782 | 0.805 (+2.94%) | 0.835 (+6.78%) |
| Subject 2 | 0.812 | 0.824 (+1.48%) | 0.855 (+5.30%) |
| Subject 3 | 0.783 | 0.795 (+1.53%) | 0.822 (+4.98%) |
| Subject 4 | 0.770 | 0.783 (+1.69%) | 0.816 (+5.97%) |
| Subject 5 | 0.839 | 0.793 (-5.48%) | 0.836 (-0.36%) |
| **Average** | **0.797** | **0.782 (+0.43%)** | **0.833 (+4.53%)** |

## Discussion and Conclusion

Spatiotemporal interpolation was used in the proposed method to correct severe motion artifact in in-utero fMRI data. Our preliminary data especially demonstrated its performance in correcting geometric mismatches between slices, as shown in the improvement in structural similarity of each volume with template. Our work offers technical advances for reliable fMRI studies in the moving fetus.

## References

1. Ferrazzi G, Kuklisova Murgasova M, Arichi T, et al. Resting State fMRI in the moving fetus: A robust framework for motion, bias field and spin history correction. Neuroimage [Internet] 2014;101:555–568. doi: 10.1016/j.neuroimage.2014.06.074.

2. Seshamani S, Blazejewska AI, Mckown S, Caucutt J, Dighe M, Gatenby C, Studholme C. Detecting default mode networks in utero by integrated 4D fMRI reconstruction and analysis. Hum. Brain Mapp. [Internet] 2016;0. doi: 10.1002/hbm.23303.

3. You W, Evangelou IE, Zun Z, Andescavage N, Limperopoulos C. Robust preprocessing for stimulus-based functional MRI of the moving fetus. J. Med. Imaging [Internet] 2016;3:26001. doi: 10.1117/1.JMI.3.2.026001.

4. You W, Serag A, Evangelou IE, Andescavage N, Limperopoulos C. Robust motion correction and outlier rejection of in vivo functional MR images of the fetal brain and placenta during maternal hyperoxia. In: Gimi B, Molthen RC, editors. Proc. SPIE 9417, Medical Imaging 2015: Biomedical Applications in Molecular, Structural, and Functional Imaging. Orlando, FL: SPIE; 2015. p. 941700. doi: 10.1117/12.2082451.

5. Jenkinson M, Bannister P, Brady M, Smith S. Improved Optimization for the Robust and Accurate Linear Registration and Motion Correction of Brain Images. Neuroimage [Internet] 2002;17:825–841. doi: 10.1006/nimg.2002.1132.

6. Klein S, Staring M, Murphy K, Viergever M a., Pluim J. elastix: A Toolbox for Intensity-Based Medical Image Registration. IEEE Trans. Med. Imaging [Internet] 2010;29:196–205. doi: 10.1109/TMI.2009.2035616.

7. Wang Z, Bovik A, Sheikh HR, Simoncelli EP. Image Quality Assessment: From Error Visibility to Structural Similarity. IEEE Trans. Image Process. [Internet] 2004;13:600–612. doi: 10.1109/TIP.2003.819861.

8. Abaci Turk E, Luo J, Torrado-carvajal A, et al. Automated ROI Extraction of Placental and Fetal Regions for 30 minutes of EPI BOLD Acquisition with Different Maternal Oxygenation Episodes. In:Proc. Intl. Soc. Mag. Reson. Med. Vol. 23; 2015. p. 639

